# Impact of genomic selection on genetic diversity in five local European cattle breeds

**DOI:** 10.1101/2025.09.08.674844

**Authors:** R. Bonifazi, T.H.E. Meuwissen, P. Croiseau, G. Restoux, S. Minéry, J.J. Windig

## Abstract

Genomic selection (GS) has revolutionised animal breeding and accelerated genetic gains in breeding programs. While GS has become common in major dairy cattle breeds, its implementation in local breeds has begun only more recently or is still in progress. However, the introduction of GS in some major breeds has also been associated with increased inbreeding rates, raising concerns about the potential effects of GS on the genetic diversity in smaller or local breeds. Our aim was to investigate the impact of GS on genetic diversity in five (small) local cattle breeds from three European countries. The five breeds evaluated were: MRY (from the Netherlands), Norwegian Red (from Norway), Abondance, Tarentaise, and Vosgienne (from France). We investigated changes in population demographic structure, as well as trends and rates of kinship and inbreeding, using both pedigree- and genomic-based measures. The population size varied depending on the breed, with Vosgienne being the smallest and Norwegian Red being the largest. The dataset included 4,645 MRY, 193,489 Norwegian Red, 16,427 Abondance, 8,882 Tarentaise, and 4,466 Vosgienne genotyped animals for more than 40,000 single-nucleotide polymorphisms. Overall, following the implementation of GS in these breeds, we observed a reduction in generation intervals for sires, fewer calves that later became sires, and, for the French breeds, a broader sire usage. Such changes were likely due to GS enabling the preselection and screening of more young bulls. Additionally, we observed a more balanced contribution of the top ten sires after the introduction of GS. Although changes in inbreeding and kinship rates occurred after the introduction of GS, there was no consistent pattern across breeds: rates increased in MRY and Tarentaise, but decreased in Norwegian Red, Abondance, and Vosgienne. Our study suggests that changes and increases in inbreeding rates may occur after the introduction of GS, although they may not be directly due to the introduction of GS *per se*, but rather due to population management strategies, such as optimal contribution selection or other breeding practices implemented at the nucleus level. Our findings emphasise the importance of monitoring changes in both genetic diversity and population demographic structure after implementing GS in local breeds, as well as adjusting breeding strategies when needed to ensure long-term sustainability.

**Interpretive Summary:** Genomic selection (GS) has transformed cattle breeding. We investigated changes in population demographic structure and genetic diversity in five European breeds after the introduction of GS. Changes in inbreeding and kinship rates were not consistent across breeds, with both increases and decreases observed. Genetic management strategies, such as optimal contribution selection, had a greater impact on maintaining genetic diversity than the introduction of GS *per se*. These findings highlight the need to monitor changes in genetic diversity and population demographic structure after the implementation of GS and, when needed, to adapt management strategies to ensure long-term sustainability.

## INTRODUCTION

The advent of genomic selection (GS) (Nejati-Javaremi et al., 1997; Meuwissen et al., 2001) had a profound impact on cattle breeding (García-Ruiz et al., 2016; Meuwissen et al., 2016). Among others, the implementation of GS in cattle breeding programs increased selection accuracy, improved the pre-selection of young bulls reducing costs associated with progeny testing, increased selection intensity, and reduced generation intervals (Schaeffer, 2006; Roos et al., 2011; Wiggans et al., 2017). These factors led to higher genetic gains for traits of interest when implementing GS compared to traditional breeding schemes (Schaeffer, 2006; Hayes et al., 2013; Meuwissen et al., 2013; García-Ruiz et al., 2016). Nowadays, GS have become common in popular cattle breeds while local breeds recently started to, or are in the process of, introducing GS (Pryce and Daetwyler, 2012; Wiggans et al., 2017; Interbull Centre, 2024). However, there has been concern on whether the higher selection intensity and faster turnover of generations brought by GS may also lead to increased mating of related individuals, especially in small populations, which would then result in faster loss of genetic diversity, increased inbreeding depression, and increased frequency of recessive deleterious mutations (Cole, 2015; Howard et al., 2017; Doekes et al., 2021). It is therefore essential to investigate and monitor the possible impact of implementing GS on the genetic diversity of local cattle breeds as these breeds may present unique features in their management strategies and breeding programs compared to more popular ones.

Compared to pedigree-based selection, GS allows the estimation of Mendelian sampling terms, enabling the exploitation of within-family variation and potentially reducing the co-selection of family members (Daetwyler et al., 2007; Lillehammer et al., 2011). However, results from previous studies across several cattle breeds showed that genetic diversity may be impacted by the introduction of GS. In Holstein, inbreeding rates increased since the introduction of GS in the USA (Forutan et al., 2018; Makanjuola et al., 2020; Guinan et al., 2023), the Netherlands (Doekes et al., 2018), Poland (Topolski and Jagusiak, 2020), France (Doublet et al., 2019), Australia (Scott et al., 2021), and Italy (Ablondi et al., 2022). Similarly, Makanjuola et al. (2020) and Sarviaho et al. (2023) reported increases in inbreeding rates after the introduction of GS in Jersey and in Finnish Ayrshire, respectively. These increases were not only for rates of inbreeding on a yearly basis, which is expected due to the shortening of generation intervals when implementing GS (Pryce and Daetwyler, 2012), but also on a generational basis. On the other hand, similar increases in inbreeding rates were not observed for other breeds where GS has also been introduced, such as Normande and Montbéliarde (Doublet et al., 2019) and Aberdeen Angus (Lozada-Soto et al., 2023). In particular, Doublet et al. (2019) showed that the breeding scheme had an impact on the effect of GS on genetic diversity. Thus, the impact of the implementation of GS on the genetic diversity of cattle breeds may differ across breeds.

In the pre-genomics era, pedigree-based inbreeding was the only tool to assess genetic diversity, genetic drift, and inbreeding. Nowadays, genomic data offers new tools to further evaluate and assess inbreeding in cattle populations throughout their genome (Howard et al., 2017; Doekes et al., 2021). The aim of this study was to investigate, using both pedigree-based and genomic-based measures, the impact of GS on genetic diversity in local cattle populations. In particular, we investigated inbreeding rates before and after the introduction of GS in five (small) local European cattle breeds from three countries. These breeds have undergone GS for at least four years and differ both in population size and genetic management, allowing to investigate the impact of GS on inbreeding rates and loss of genetic diversity relative to each population’s demographic structure (e.g., number of sires/dams, sires’ contribution, etc.), which can vary significantly across local breeds.

## MATERIAL AND METHODS

The data analysed in this study came from existing databases and was provided by the respective breed organisations. Thus, no animals were used in this study, and ethical approval for the use of animals was not necessary.

### Data available

Individual pedigree and genomic data were available for five local European cattle breeds across three countries: Meuse Rhine Yssel (MRY) from the Netherlands, Norwegian Red Cattle (NRC) from Norway, and Abondance (ABO), Tarentaise (TAR), and Vosgienne (VOS) from France (Figure 1). Table 1 reports an overview of the size of the datasets used for each breed, the years analysed, and the year in which GS was introduced. We defined the year of the introduction of GS as the year in which genomic-based estimated breeding values were first made available to breeders. Hereafter, we provide a brief description of the breeds and describe the data preparation of each dataset.

**Figure 1.**
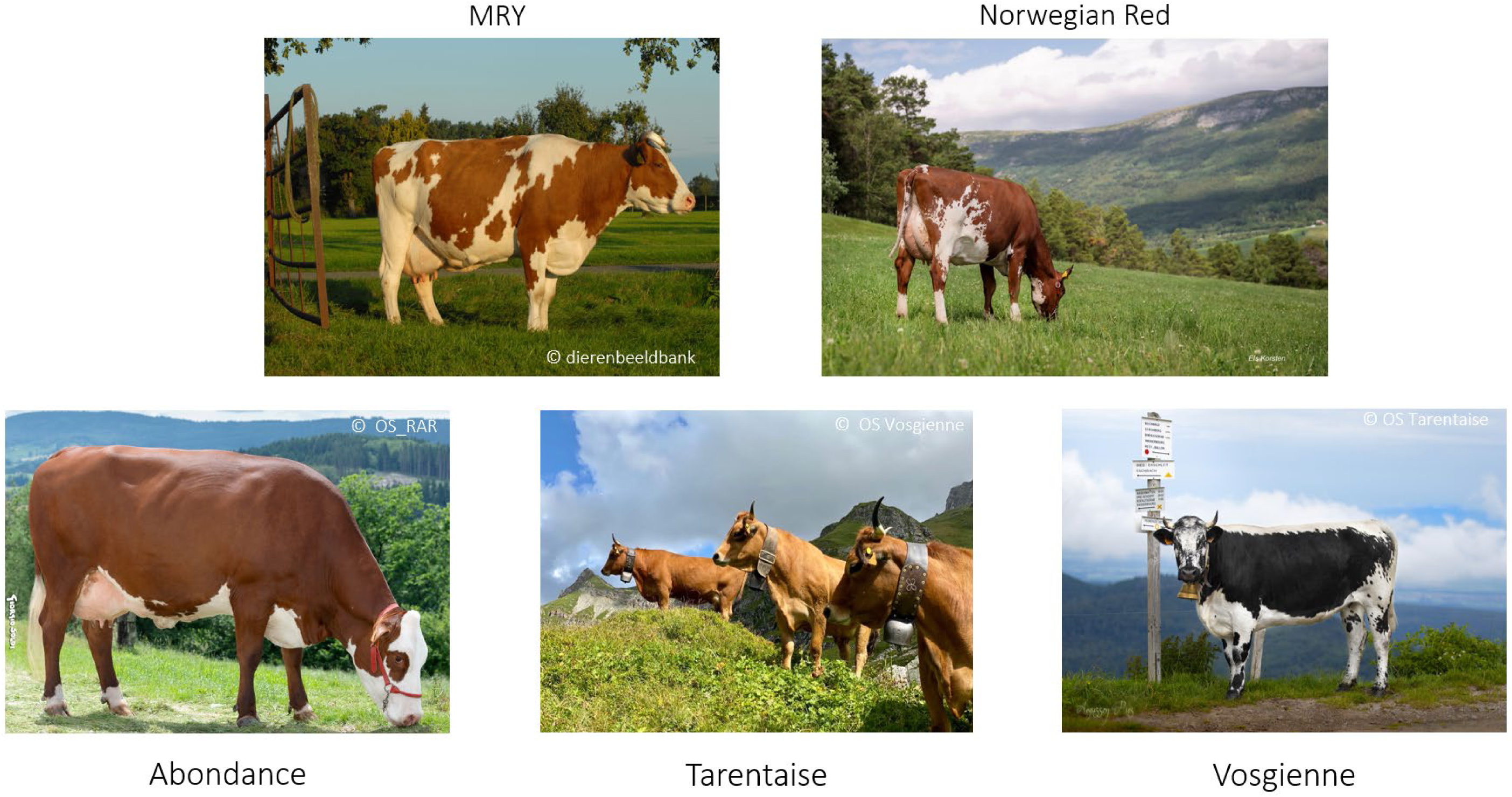
Breeds investigated in this study: MRY from the Netherlands (© dierenbeeldbank), Norwegian Red from Norway (© Els Korsten), Abondance (© OS_RAR), Tarentaise (© OS Tarentaise), and Vosgienne (© OS Vosgienne) from France.

**Table 1.** Summary of data and information available in each breed: size of the pedigree, number of genotyped animals, interval of years analysed, year of introduction of genomic selection, and number of registered breeding females in 2021/2022 (source DAD-IS FAO, 2025).

| Breed | Pedigree | Genotyped animals | Years analysed | Introduction of genomic selection | Breeding females |
| --- | --- | --- | --- | --- | --- |
| MRY | 205,933 | 4,645 | 1975-2021 | 2018 | 8,068 |
| Norwegian Red | 5,125,148 | 193,489 | 1975-2022 | 2016 | 175,975 |
| Abondance | 833,109 | 16,427 | 2000-2020 | 2014 | 44,1480 |
| Tarentaise | 279,333 | 8,882 | 2000-2020 | 2014 | 14,105 |
| Vosgienne | 101,490 | 4,466 | 2000-2020 | 2014 | 11,823 |

### Maas Rijn Yssel breed

The MRY is a Dutch dual-purpose cattle breed that originates from the central region of the Netherlands, between the rivers Meuse, Rhine, and Yssel from which it is named after. The MRY is a relatively small population (Hiemstra et al., 2010; Stoop et al., 2017; Eynard et al., 2018b) with around 5,000 calves born per year, and in which GS began in the late 2018 (CRV). The genetic management of the MRY breed partially followed that of the Dutch Holstein-Friesian breed (Doekes et al., 2018; Bonifazi et al., 2022), with the breeding goal expanded around 2000 to include traits related to fertility, reproduction, and longevity. Moreover, around the year 2000 the so-called “cold sire system” was introduced in MRY to maintain genetic diversity by increasing the number of sires used (Hiemstra and de Haas, 2004). In this system, around 10 young bulls are progeny tested per year, and each is culled one 20,000 doses of frozen semen have been collected. Although optimum contribution selection (OCS; Meuwissen, 1997) has not been formally implemented in MRY, some of its principles have been applied through the cold sire system and by ensuring that the progeny-tested bulls selected each year originated from different parents, thereby limiting coancestry among sires and restricting the rate of inbreeding (Henk Geertsema, personal communication). Pedigree and genomic data were provided by CRV (https://crv4all.com). The pedigree included 205,933 animals born between 1910 and 2022 and was checked using the “OptiSel” R package (Wellmann, 2019) for absence of duplicates, pedigree cycles, and inconsistencies in the sex of the parents. All animals in the pedigree had information on the year of birth. Before 1975, data was incomplete and results on these years are not presented, although the full pedigree information were used for calculation of inbreeding and kinship coefficients. Genomic data was available in the form of individual SNP genotypes imputed at 44,645 autosomal markers for 4,645 purebred MRY animals (i.e., with a pedigree-based breed composition for MRY ≥87.5%). The SNP panel was checked for the absence of duplicated SNPs (one duplicated SNP removed). Further quality controls on genomic data were performed using Plink v1.9 (Chang et al., 2015): SNPs with minor allele frequency lower than 0.01 were discarded (3,685 SNPs removed). After the above edits, 4,645 animals and 40,959 SNP remained for subsequent analyses (Table 1).

### Norwegian Red breed

The Norwegian Red (NRC) cattle breed, locally known as Norsk Rødt Fe, originated in Norway around 1930. The pedigree included 5,125,148 animals, born between 1900 and 2024. Mating decisions were based on the relationships between the parents and their estimated breeding values. The overall inbreeding was monitored and when needed reduced by selecting more sires with lower pedigree relationships. From 2010 onwards, the genetic management has increasingly been using the mating proportions recommended by OCS. Before genomic selection, about 100 young bulls were progeny tested annually, of which about 15 were selected to be AI sires. Assuming a sex ratio of 0.5 the number of calves born per year is around 100,000 per year and NRC was the largest breed analysed in this study (Table 1). Individual genotypes were available for a panel of 45,255 SNPs on 193,489 animals (MAF > 0.01). For NRC, GS started in 2016. Finally, results for NRC are presented up to and including 2022 for consistency between results based on genomic data (available until 2022) and those based on pedigree.

### Abondance, Tarentaise, Vosgienne

The Abondance (ABO), Tarentaise (TAR), and Vosgienne (VOS) are all French cattle breeds adapted to mountain conditions. These local breeds are essential to the preservation of mountain regions and are linked to the production of numerous PDO (Protected Designation of Origin) French cheeses, making them profitable despite their small population sizes.

ABO, originally from Haute-Savoie, is a dairy breed renowned for its milk, which is used in the production of cheeses such as Reblochon or Abondance. ABO is the fourth French dairy cattle breed in terms of population size with about 43,000 live cows (Institut de l’Élevage, 2022), albeit it represents only 1.3% of the total French dairy cattle population. With over 25,000 calves born per year it was the second largest breed analysed in this study. The breed originates from the French Alps where it usually grazes on pasture, and it is well adapted to high altitudes (>2,000 meters). The ABO breed is mainly a free-range breed even if it is raised indoor during the winter season due to the rigorous conditions in the mountains. Cows produce about 6,000 kg of milk per year. The mean herd size of this breed is about 20 cows. Reproduction is almost exclusively conducted with artificial insemination (AI). The selection nucleus consists of 1,261 farms and 21,000 cows (equivalent to 38% of the total population). The pedigree included 833,109 animals born between 1946 and 2021. Each year about 20 AI sires are selected and GS has started in 2014. Finally, ABO was the largest among the French breeds analysed (Table 1).

TAR is an excellent dairy breed also originating and grazing in the Alps, and whose rich milk is used to make alpine cheeses, such as Beaufort. It is often found in the same area as ABO and raised under the same conditions: mainly free-range, except during winter. As for ABO, feeding mainly relies on hay when indoor and silage is usually not used. It produces about 4,800 kg of milk per year per cow. Its live population size consists of about 14,000 cows (Institut de l’Élevage, 2022), and with around 9,500 calves born per year, its population size is intermediate among the five breeds analysed in this study. ABO has a selection nucleus of 424 farms including 7,700 cows (equivalent to 59% of the total population). Each year 18 AI sires are selected. The TAR pedigree included 279,333 animals born between 1949 and 2021.

VOS, originally from the Vosges mountain area, is a dual-purpose breed, well known for its milk (used in the production of Munster cheese) and for its ability to efficiently utilize poor-quality and marginal pastures. VOS is selected for both milk and meat, with cows producing about 4,200 kg of milk per year. The VOS consists of a small French population located in the eastern Vosges massif. With less than 4,000 calves born per year, VOS is the smallest breed analysed in this study. The pedigree included a total of 101,490 animals born between 1950 and 2021. Like the other mountainous local French breeds, it is raised free-range except during winter. The population consists of about 10,000 cows in total, with only 4 to 6 AI sires selected each year (Effectifs et localisation - La Race Bovine Vosgienne, OS Vosgienne personal communication). The selection nucleus consists of 79 farms and 1,400 cows (equivalent to 15% of the total population). In VOS, all animals are genotyped, making it the only breed in France to fall into such situation.

For the three French breeds, individual genotypes were available at 50K SNPs, or imputed to 50K. For the French breeds, 53,459 SNPs were initially available. A filter was applied to remove monomorphic or absent variants within each breed, leading to a remaining 48,178 SNPs in ABO, 46,109 SNPs in TAR, and 45,747 SNPs in VOS. The resulting call-rate was 100% in every breed. Only genotypes of animals born from 2012 onwards were considered to get a representative picture of the whole population since from this date a large majority of the animals were genotyped in each of the three French breeds. The number of genotyped animals considered were 16,427, 8,882, and 4,466 for ABO, TAR, and VOS, respectively (Table 1). Finally, GS started in 2014 in all three French breeds.

### Population demographic structure and statistics

Population demographic structure and other related statistics were obtained based on pedigree data using Retriever software (Windig and Hulsegge, 2021). The average generation interval (*L*) per year, defined as the average age of the parent(s) at birth of their offspring, was computed for sires, dams, and both parents. To quantify the size of the population involved in breeding and its changes over time, we computed for each year: the number of born male and female calves which later have become sires and dams, respectively; and the number of bulls that sired calves in that year; and the average number of calves born per sire. Finally, to investigate the skewness in sires’ contributions, an important cause of high inbreeding rates, we quantified changes in the contribution of the ten most used sires within a specific year to the total number of offspring born in that same year. Top ten sires’ contributions were expressed as percentages.

### Pedigree-based and genomic-based measures of genetic diversity

Pedigree-based measures of genetic diversity were computed using Retriever software (Windig and Hulsegge, 2021). These measures were calculated using the whole pedigree and averaged per year. Inbreeding coefficients for each animal in the pedigree (*F_PED_*) were calculated following Meuwissen and Luo (1992). Kinship coefficients (*f_PED_*) between all animals born in the same year were calculated following Sargolzai et al. (2005). Average *f_PED_* were computed without self-kinship for all animals in the pedigree, and for animals later becoming sire or dam.

For genotyped animals, genomic inbreeding (*F_ROH_*) was calculated based on runs of homozygosity (ROHs) detected using Plink v1.9 (Chang et al., 2015) and analysed using the “detectRUNS” R package (Biscarini F et al., 2019). The following Plink parameters were used to estimate ROHs: *--homozyg-window-snp 15, --homozyg-gap 150, --homozyg-snp 15, -- homozyg-density 75, --homozyg-kb 1000*. Thus, ROHs were defined using a scanning window of 15 consecutive homozygotes SNP. ROHs must have had a maximum gap between two SNPs of 150 kb, a minimum of 15 SNP, a minimum of one SNP per 75 kb, and a minimum length of 1,000 kb. Then, the *F_ROH_* for individual *i* was calculated as the proportion of genome in ROH (McQuillan et al., 2008) as 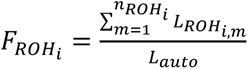, where 𝑛*_ROH_i* is the total number of ROHs in individual *i*, 𝐿*_ROHi,m_* is the length of the *m^th^* ROH, and 𝐿_*auto*_ is the total length of the autosome covered by SNPs.

### Rates of inbreeding

Annual and generation rates of inbreeding were computed for each breed for specific intervals of years based on the observed major breakpoints in pedigree-based inbreeding trends and the year of introduction of GS. For MRY, these periods were 1975-2000, 2000-2013, 2013-2019, and 2019-2021. For NRC, the defined periods were 1975-2000, 2000-2010, 2010-2016, and 2016-2020. Finally, for ABO, TAR, and VOS the defined periods were 2000-2014, and 2014-2020, corresponding to progeny testing and genomic evaluations periods, respectively. Annual rates of inbreeding for *F_PED_* and *F_ROH_* coefficients were computed following Pérez-Enciso (1995), i.e., as −1 times the slope of the regression of 𝑙𝑛(1 − 𝑥̅) on the year of birth, with 𝑥𝑥̅ being the average value of either *Fped* or *Froh* per year. This formula accounts for the fact that inbreeding is not linear and asymptotes to 1. Finally, inbreeding rates per generation were obtained by multiplying the annual rates of inbreeding by the average generation interval of both parents in the years considered. Inbreeding rates were expressed as percentages per year or per generation (*ΔF*).

### Software availability

The Retriever software (Windig and Hulsegge, 2021) is accessible at https://genebankdata.cgn.wur.nl/software/Retriever/Retriever.html. R functions used to process the outputs of Retriever have been made available in the “labradoR” R package, available at: https://github.com/bonifazi/labradoR.

## RESULTS

Hereafter, we first present the results on the population demographic structure for the five breeds, followed by the results on genetic diversity measures.

### Population demographic structure Population size

Across all breeds, the number of male and female calves that later became sires and dams, respectively, fluctuated across years but remained overall rather constant, with the exception of MRY and NRC (Figure 2). For MRY, the number of calves that later became sire born per year constantly decreased over time: from 367 in 1975 to 85 in 2018 (Figure 2). On the other hand, the number of MRY calves that later become dams remained stable up to 2016, fluctuating around 1,500 per year (1,337 and 1,439 in 1975 and 2016, respectively), to then decrease afterwards (1,046 in 2018). For NRC, the number of calves that later become sires was stable at around 400 per year between 1975 up to 2010 (365 and 656, respectively), with the exception of a sharp increase at around the years 2000 and 2001 (816 in 2000). From 2009 onwards, the number of calves that later become sires increased again at around 600 per year (654 in 2010) (Figure 2). Overall, the number of NRC calves that later become dams fluctuated around 60,000 per year (58,660, 65,001, and 58,443, in 1975, 1981, and 2016, respectively). For the French breeds, the number of calves that later become sires remained about the same up to and after 2014: on average across all years 148.6, 52.5, and 17.5, for ABO and TAR, and VOS, respectively. The number of calves that later become dams increased for all French breeds from 2000 to 2014 or 2015, and slightly decreased afterwards: from 5,898, 2,135, 390 in 2000, to 7,417, 2,770 and 626 in 2014, and 641, 342, and 58 in 2018, for ABO, TAR, and VOS, respectively. A slight decrease in the most recent years of the number of male and female calves that later become becoming sires and dams, respectively, is expected because animals may still be too young to have become parents yet.

**Figure 2.**
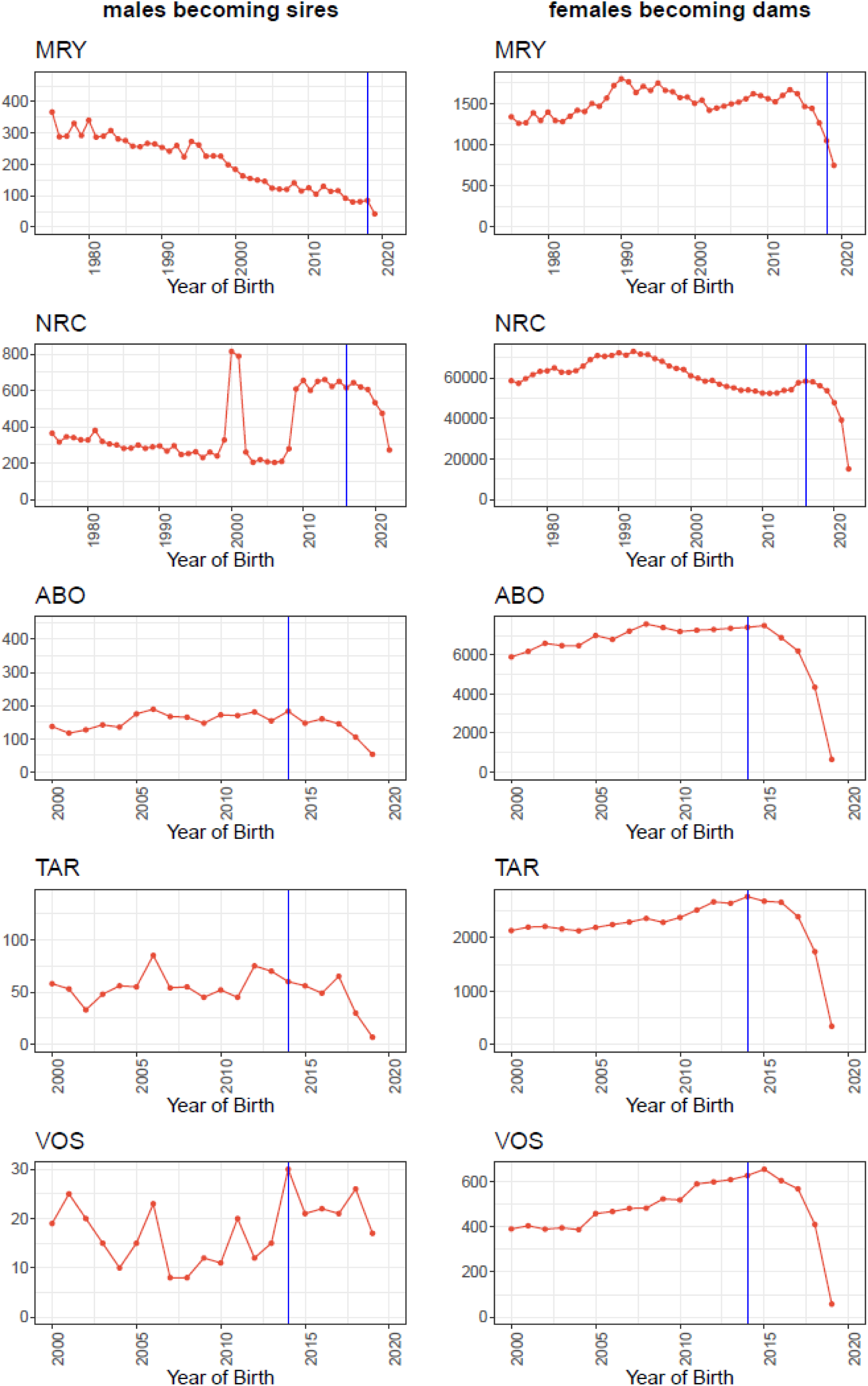
Number of males becoming sires and of females becoming dams in each year of birth. Vertical blue horizontal lines indicate the year of onset of genomic selection. The last three (for MRY) or two (other breeds) years were removed as males and females are still young to be used as parents.

The number of sires used per year decreased over time for MRY, while it increased for NRC and French breeds. For MRY, the number of sires used in each year decreased from 607 in 1975 to 362 in 2005 (Figure 3), remaining overall stable in the years before and after the implementation of GS (on average 360.3 between 2005 and 2021). For NRC, the number of sires used in each year gradually decreased from 1975 to 2000 (908 and 547, respectively). From 2000, the number of sires used increased up to 2015 (1,209 sires), with two sharp increases in 2001-2002 and in 2011. These increases are due to a more detailed registration of natural mating NRC sires on commercial farms. However, after the implementation of GS, the number of sires used in NRC started to decrease, moving from 1,150 in 2016 to 872 in 2022 (Figure 3). This decline is attributed to a decrease in the number of natural mating sires, following a general trend towards reduced use of such sires. Although natural mating sires dominate the number of sires that have offspring, their total number of offspring is small compared to that of AI sires, and their offspring are rarely selected as parents, i.e. their contribution to the genetic improvement of the NRC population is small. The recent reduction in the use of natural mating sires has been accompanied by a corresponding increase in the number of calves born per sire (Supplementary Figure S1). Finally, for all three French breeds, the number of sires used each year increased over time, even after the introduction of GS (Figure 3): from 288, 122, and 44 in 2000 to 437, 177, and 69 in 2014, and to 449, 168, and 100 in 2020, for ABO, TAR, and VOS, respectively. Thus, after the introduction of GS, the number of sires used remained stable (MRY), decreased (NRC), or increased (ABO, TAR, VOS).

**Figure 3.**
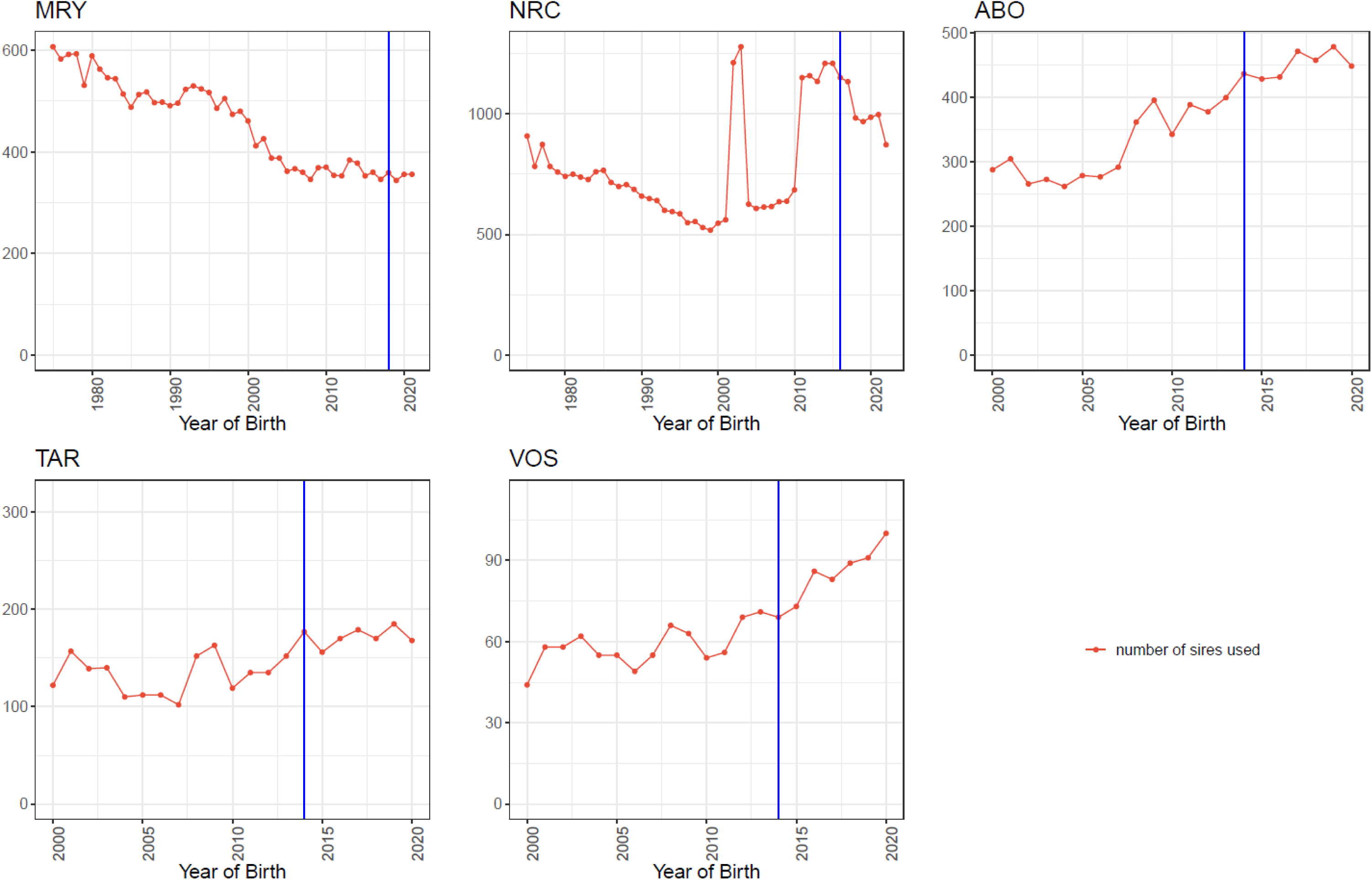
Number of sires used in each year. Vertical blue horizontal lines indicate the year of onset of genomic selection.

### Generation interval

Overall, the average age of dams at time of calving was more or less constant over time in all five breeds, even after the implementation of GS (Figure 4). In general, the dams’ generation interval was close to 5 years or slightly above for all breeds except for the NRC, for which it was around 3.5 years. Compared to dams, generation intervals for sires were less stable (Figure 4). For MRY, sires’ generation intervals fluctuated over time and was on average 6.8 years across the whole period 1975-2021, showing a small decrease before the implementation of GS to then slightly increase afterwards (on average 7.8, 6.7, 6.3, and 6.6 in 1975-1980, 1981-2015, 2015-2018, and 2019-2020). For NRC, the sire generation interval was on average 5.5 years between 1975 up to the introduction of GS in 2016, after which it strongly decreased to close to 3 years from 2018 onwards (average of 2.9 years between 2018 and 2022). For ABO, the sires’ generation interval slightly increased up to the introduction of GS (7.4 to 8.9 years in 2000 and 2014, respectively), after which it decreased to 4.9 years in 2020. For TAR, the sires’ generation interval fluctuated before the introduction of GS, with values above 7.5 years (maximum of 8.5 in 2010), and decreased afterwards, up to 4.2 years in 2020. In VOS, sires’ generation interval increased from 7.4 years in 2000 to 10.4 years in 2015, remaining at this high level for three years after the introduction of GS, to then decrease to 8.5 years in 2020. Thus, sires’ generation intervals clearly decreased in all breeds after the introduction of GS, except for the MRY.

**Figure 4.**
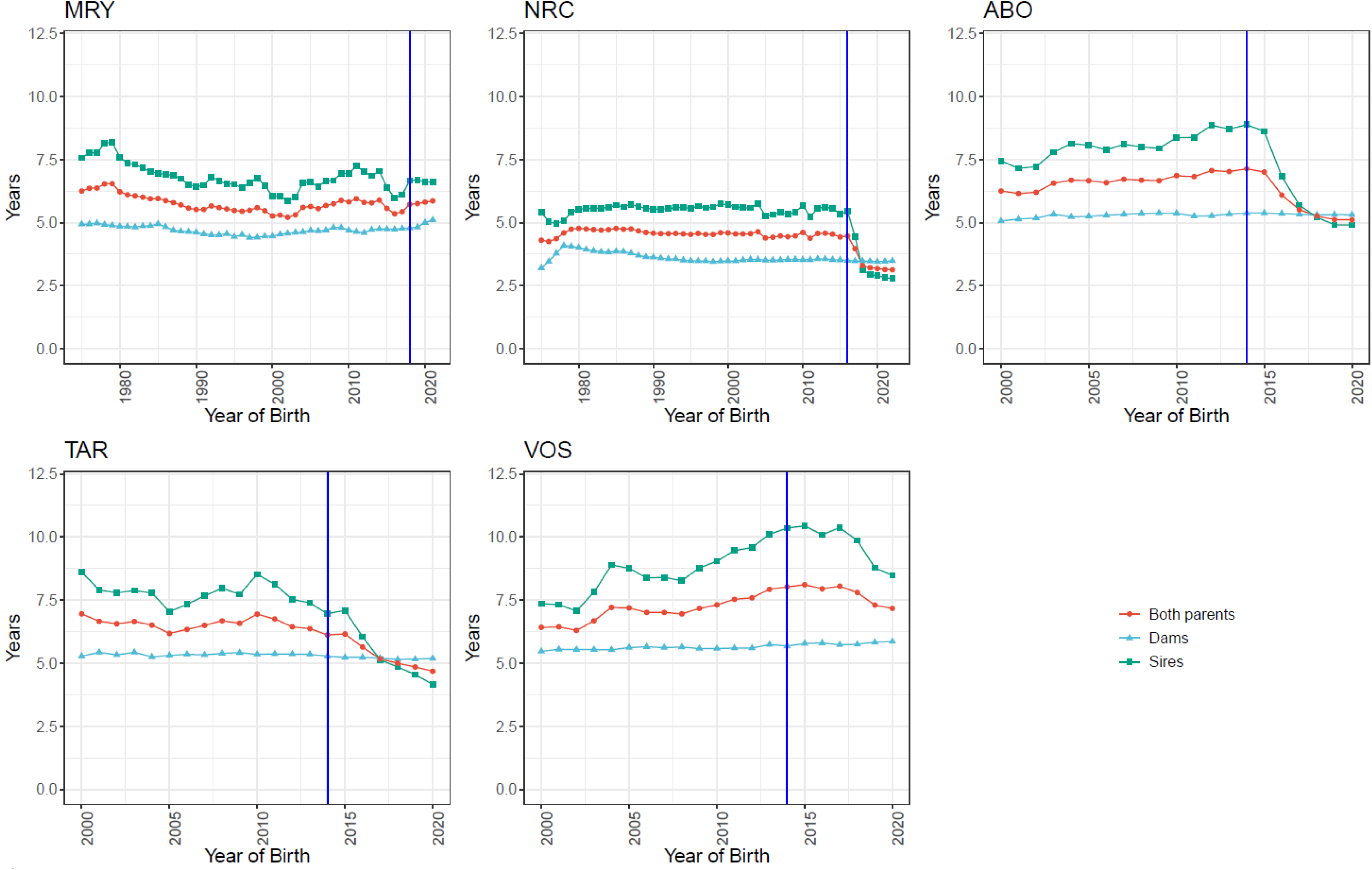
Generation intervals trends for sires, dams, and both parents. Y-axes report the number of years. Vertical blue horizontal lines indicate the year of onset of genomic selection.

### Skewness of sires’ contributions

For all breeds, the contribution of the top 10 sires reduced in recent years (Figure 5). For MRY, the contribution of the top 10 MRY sires increased from 28.4% in 1975 to over 46.1% in 1996 (Figure 5). From 1997 to 2013, the contribution of the top 10 sires varied between 35.5% and 43.8%, then reduced to an average of 35.5% between 2014 and 2018, and of 33.9% in the years after the implementation of GS (2019-2021). For NRC, the contribution of the top 10 sires increased over time from 17.9% in 1975 to 52.3% in 2014 (Figure 5), to then decrease, especially in the years after the introduction of GS, down to 28.8% in 2022. For both MRY and NRC, the contributions within the top 10 sires were quite even in the last years, while in the years around 2000, the contributions were far more skewed, with the top 3 sires taking close to or more than half of the top 10 sires’ contributions (51.7% and 43.5% on average for the period 2000-2015, for MRY and NRC, respectively). Overall, French breeds showed an high contribution of the top 10 sires, especially in the years before the implementation of GS. Despite being the largest of the French breed analysed, the contribution of the top 10 sires in ABO were above 50% up to 2015 (54.8% on average for the period 2000-2015; Figure 5). After 2015, i.e., a year after the introduction of GS, the contribution of the top 10 sires strongly reduced to 29.6% on average for the period 2016-2020. For TAR, the pattern was similar to ABO; the contribution of the top 10 sires was 50.9% on average for the period 2000-2015, and then reduced to 31.9% for the period 2016-2020. Among all breeds, VOS had the highest contribution of the top 10 sires, with values above 65% and close to 75% in the period 2000-2011 (average of 70.1% and 68.0% for the period 2000-2011 and 2000-2014, respectively). Such high contributions for VOS were expected given the limited population size and number of selected AI sires per year. After 2012, the contribution of the top 10 sires in VOS started to reduce and was below 50% from 2018 onwards (51.8% and 48.9% on average for the period 2014-2020 and 2016-2020, respectively). Overall, the observed reduction in all breeds of the contribution of the top 10 sires happened either before (MRY) or after (NRC and French breeds) the start of GS.

**Figure 5.**
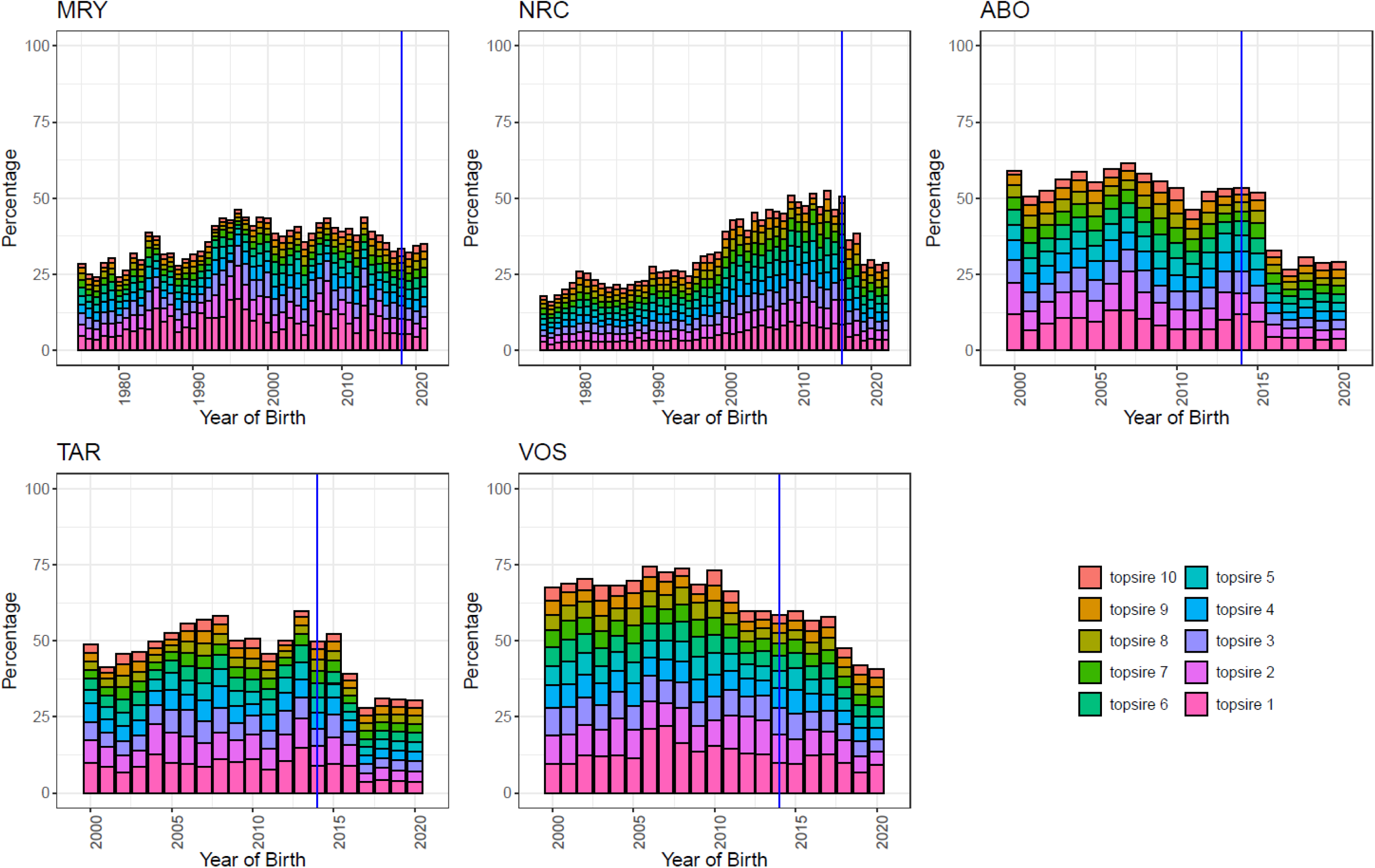
Contribution (in percentage) of the top ten sires for animals born in each year. Vertical blue horizontal lines indicate the year of onset of genomic selection.

### Genetic diversity

Trends of pedigree-based inbreeding and their corresponding annual and generational rates (Δ*F_PED_*) varied according to the specific breed considered. For MRY, *F_PED_* increased from 1975 up to 2013, decreased after 2013 up to 2019, and finally increased again in 2020 and 2021 (Figure 6). Before the introduction of GS, rates of *F_PED_* were always positive both on an annual and on a generational base, except for the period 2013-2019 (Table 2): *F_PED_* of 0.10%/year, 0.08%/year, and −0.12%/year, and Δ*F_PED_* of 0.61%, 0.46%, and −0.67%, for the periods 1975-2000, 2000-2013, and 2013-2019, respectively. After the introduction of GS, rates of pedigree-based inbreeding were again positive: *F_PED_* of 0.09%/year and Δ*F_PED_* of 0.51% for the period 2019-2021. For NRC, *F_PED_* increased from 1975 until the introduction of GS in 2016 (Figure 6). Before the introduction of GS, annual rates of *F_PED_* ranged between 0.04%/year and 0.06%/year (Table 2). Similarly, Δ*F_PED_* increased from 0.24% for the period 1975-2000 to 0.26% for the period 2010-2016, with a smaller *F_PED_* for the period 2000-2010 (Δ*F_PED_* of 0.16%) (Table 2). After the introduction of GS, *F_PED_* kept increasing (Figure 6), although with a much lower rate: 0.02%/yr for the period 2016-2020, with a corresponding Δ*F_PED_* of 0.05% (Table 2). For ABO, *F_PED_* increased steadily for the whole 2000-2020 period (Figure 6). Before the implementation of GS, the annual rate of *F_PED_* was 0.18%/year for the period 2000-2014, with a corresponding Δ*F_PED_* of 1.19%. After the introduction of GS, *F_PED_* kept increasing although with a lower generational rate due the shorter generation interval: annual rate of *F_PED_* was 0.17%/year while Δ*F_PED_* was 0.99% for the period 2014-2020 (Table 2). For TAR, *F_PED_* was rather stable until 2008, then increased over time, also after the introduction of GS (Figure 6). Both annual and generational rates of pedigree-based inbreeding increased after the introduction of GS (Table 2): Δ*F_PED_* was 0.35% in 2000-2014 and 0.93% in 2014-2020 (Table 2). VOS had the lowest level of *F_PED_* among all French breeds but increased steadily over time (Figure 6). However, after the introduction of GS, both annual and generational rates of *F_PED_* decreased, with Δ*F_PED_* moving from 0.53% for the period 2000-2014 to 0.23% for the period 2014-2020 (Table 2). Thus, following the introduction of GS, Δ*F_PED_* increased in some breeds (MRY and TAR) , while it decreased in others (NRC, VOS, and slightly for ABO).

**Figure 6.**
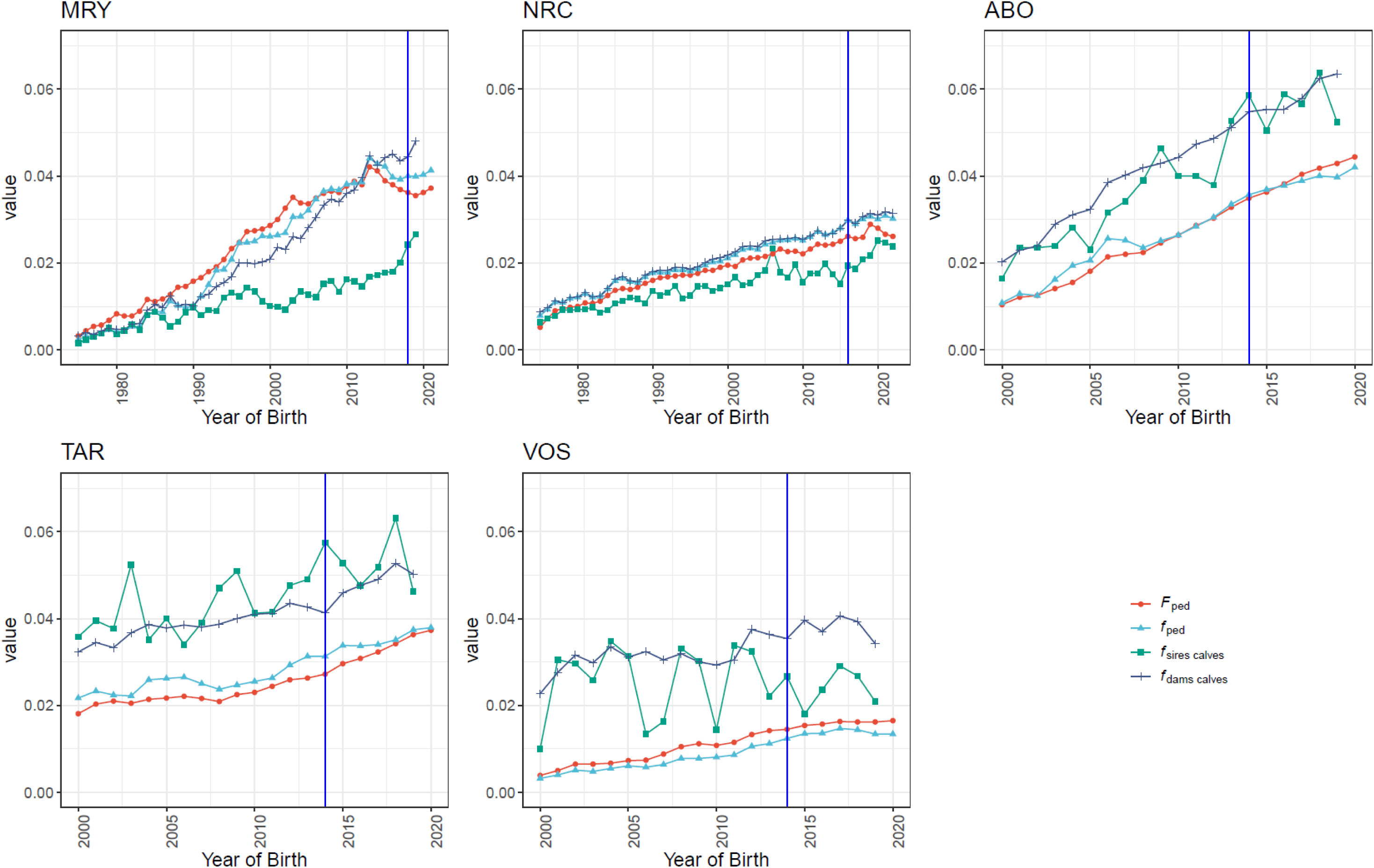
Yearly average of pedigree-based coefficients of inbreeding (*F_ped_*) and kinship for all animals in the pedigree (*f_ped_*), for calves later selected as sires (*f_sires calves_*) and for calves later selected as dams (*f_dams calves_*). Vertical blue horizontal lines indicate the year of onset of genomic selection. Kinship measures for dams and sires in years 2020-2021 and 2021 were removed for MRY and French breeds, respectively, as no parents were available in these years yet.

**Table 2.** Average generation interval for both parents (*L*) in different periods (Years Interval) for all animals in pedigree (all) and for genotyped animals only (genotyped), along with annual and generation (*Δ*) rates of pedigree-based kinship (*f_PED_*) and inbreeding (*F_PED_*) for all animals in the pedigree and of genomic-based inbreeding (*F_ROH_*) for genotyped animals only.

| Breed | Years Interval | *L* |  | *Annual*<sup>1</sup> |  |  | *Generation ($\Delta$)*<sup>1</sup> |  |  |
| --- | --- | --- | --- | --- | --- | --- | --- | --- | --- |
| | | all | genotyped | $f_{PED}$ | $F_{PED}$ | $F_{ROH}$ | $f_{PED}$ | $F_{PED}$ | $F_{ROH}$ |
| MRY | 1975-2000 | 5.84 | 6.85 | 0.10 | 0.10 | 0.11 | 0.61 | 0.61 | 0.76 |
|  | 2000-2013 | 5.61 | 6.21 | 0.13 | 0.08 | 0.08 | 0.73 | 0.46 | 0.47 |
|  | 2013-2019 | 5.64 | 5.28 | -0.08 | -0.12 | -0.21 | -0.45 | -0.67 | -1.12 |
|  | 2019-2021 | 5.81 | 5.40 | 0.07 | 0.09 | 0.17 | 0.42 | 0.51 | 0.93 |
| NRC | 1975-2000 | 4.60 | 4.79 | 0.05 | 0.05 | 0.02 | 0.22 | 0.24 | 0.09 |
|  | 2000-2010 | 4.52 | 4.65 | 0.04 | 0.04 | 0.04 | 0.20 | 0.16 | 0.18 |
|  | 2010-2016 | 4.51 | 4.55 | 0.06 | 0.06 | 0.02 | 0.29 | 0.26 | 0.10 |
|  | 2016-2022 | 3.48 | 3.37 | 0.02 | 0.02 | 0.02 | 0.06 | 0.05 | 0.06 |
| ABO | 2000-2014 | 6.67 | 6.59 | 0.17 | 0.18 | 0.42 | 1.13 | 1.19 | *2.76 |
|  | 2014-2020 | 5.89 | 5.87 | 0.10 | 0.17 | 0.12 | 0.58 | 0.99 | 0.68 |
| TAR | 2000-2014 | 6.55 | 6.43 | 0.06 | 0.05 | 0.10 | 0.38 | 0.35 | *0.64 |
|  | 2014-2020 | 5.37 | 5.39 | 0.11 | 0.17 | 0.18 | 0.56 | 0.93 | 0.96 |
| VOS | 2000-2014 | 7.12 | 6.98 | 0.06 | 0.07 | 0.07 | 0.42 | 0.53 | *0.51 |
|  | 2014-2020 | 7.77 | 7.61 | 0.01 | 0.03 | 0.01 | 0.10 | 0.23 | 0.09 |

<sup>1</sup> rates are expressed in percentage. \* coefficients were calculated for the interval 2012-2014 using available genomic data.

Overall, trends and rates of kinship for all animals in the pedigree followed those of pedigree-based inbreeding, particularly in relation to changes before and after the implementation of GS. For MRY, kinship levels (*f_PED_*) were smaller than *F_PED_* until 2007, showing avoidance of inbreeding mating. Levels of *f_PED_* were then about equal to those of *F_PED_* up to 2013, and larger for the period 2013-2021 (Figure 6). Kinships differed clearly between sires and dams after 1980 (Figure 6), with kinship between sires being considerably smaller than both kinships between dams and *F_PED_* (Figure 6), which indicates that selected sires were recruited from animals less related than the population average. Overall, for MRY, generational rates of kinship (Δ*f_PED_*) were similar to Δ*F_PED_*, being negative before the implementation of GS (-0.45%) for the period 2013-2019, and positive afterwards (0.42%) for the period 2019-2021), although with a smaller rate than before 2013 (0.73%) (Table 2). For NRC, *f_PED_* was slightly higher than F*_PED_*, but overall followed a similar trend (Figure 6). Kinships between dams were clearly higher than those between sires, with *f_PED_* for these latter increasing more after 2016. These lower kinships for sires indicates that male calves later becoming sires were selected from animals less related than average. Finally, rates of kinships in NRC were the same (annual rates) or similar (Δ*f_PED_*) to those of pedigree-based inbreeding (Table 2), with a smaller Δ*f_PED_* after the introduction of GS. For the French breeds, *f_PED_* followed overall the same trend as for *F_PED_* (Figure 6). In ABO, *f_PED_* was the same or slightly higher than *F_PED_* until 2010, to then become slightly lower after the implementation of GS. In TAR, *f_PED_* was always higher than *F_PED_*, while in VOS it was the opposite (Figure 6). In all three French breeds, kinships between sires and dams were clearly higher than *f_PED_* of all animals, indicating that parents were recruited from animals that were more related than the population average. Finally, after the implementation of GS, Δ*f_PED_* increased slightly for TAR, while it decreased for ABO and VOS (Table 2). Thus, after the introduction of GS, Δ*f_PED_* increased for MRY and TAR, while it decreased for NRC, ABO, and VOS.

Overall, genomic-based inbreeding rates followed a similar pattern as pedigree-based ones. For MRY, *F_ROH_* increased steadily from 0.054 in 1975 up to 0.083 in 1998, remaining stable around 0.08 until 2013 (Figure 7). Similar to *F_PED_*, *F_ROH_* showed a decreasing trend for the period 2013-2019 (*F_ROH_* of 0.069 in 2019), and a slight increase from 2019 onwards (Figure 7). Before GS, the generational rates of genomic-based inbreeding (Δ*F_ROH_*) were positive except for the period 2013-2019, and increased again to similar levels after the introduction of GS (Δ*F_ROH_* of −1.12% and 0.93% in 2013-2019 and 2019-2021, respectively; Table 2). For NRC, *F_ROH_* was overall stable across years (Figure 7). Annual and generational rates of genomic-based inbreeding in NRC were overall smaller than pedigree-based ones (Table 2), and decreased in the period before the introduction of GS: Δ*F_ROH_* of 0.18, 0.10, and 0.06 for the period 2000-2010, 2010-2016, and 2016-2022, respectively (Table 2). For the French breeds, only three years of genomic data were available for the period before the implementation of GS. For ABO, Δ*F_ROH_* was much higher than Δ*F_PED_*: 2.76% for the period 2012-2014 and decreased after the introduction of GS (0.68% in 2014-2020; Table 2). In TAR, Δ*F_ROH_* increased after the introduction of GS from 0.64% for the period 2011-2014 to 0.96% for the period 2014-2020 (Table 2). For VOS, Δ*F_ROH_* decreased after the introduction of GS from 0.51% for the period 2011-2014 to 0.09% for the period 2014-2020 (Table 2). Thus, after the implementation of GS, Δ*F_ROH_* increased for MRY and TAR, but decreased in NRC, ABO, and VOS.

**Figure 7.**
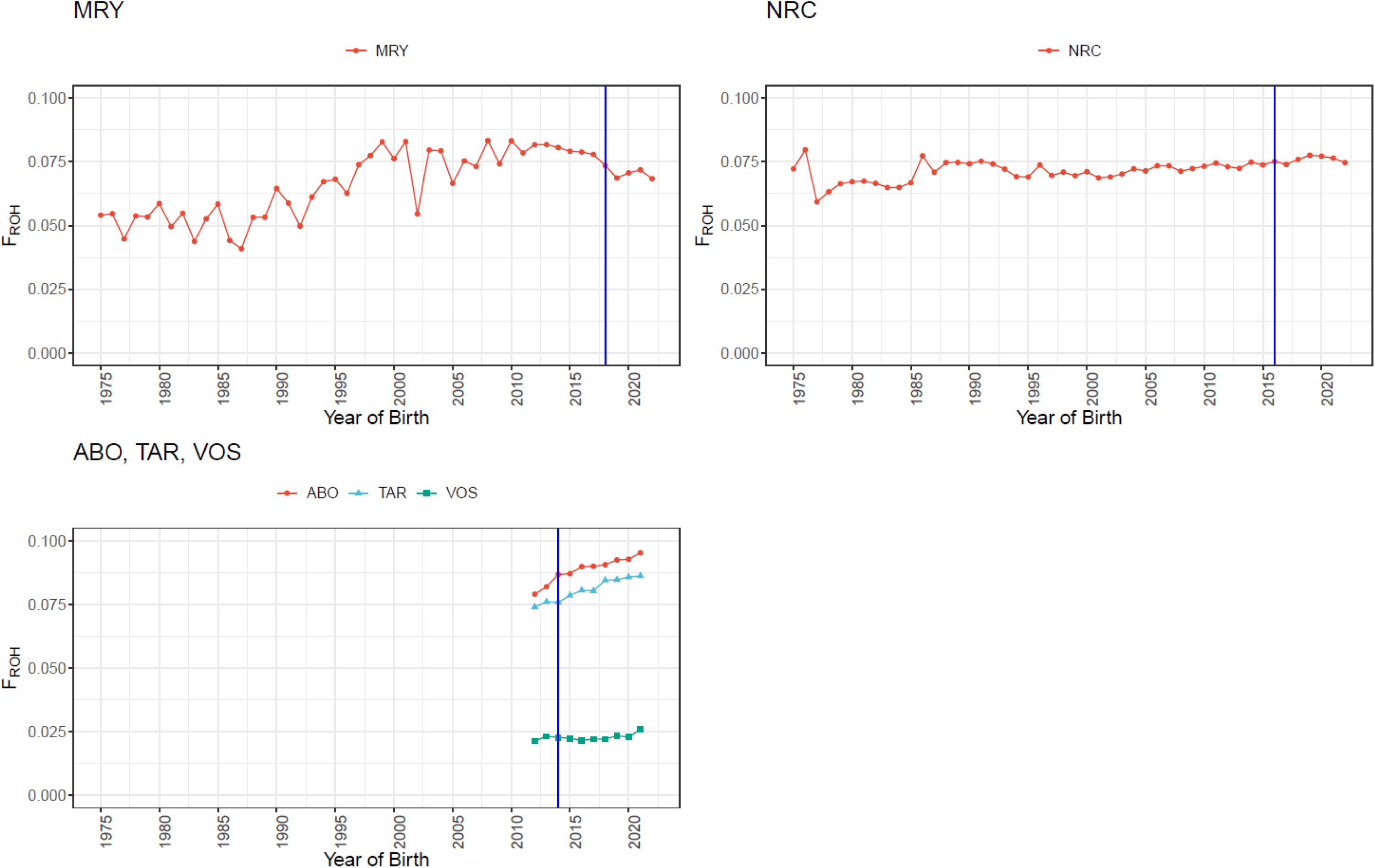
Trends of genomic inbreeding (*F_ROH_*) for genotyped animals across different breeds. Vertical blue horizontal lines indicate the year of onset of genomic selection.

## DISCUSSION

Genomic selection has had a profound influence on dairy cattle breeding leading to changes in the structure of breeding programs (Schaeffer, 2006; Van Eenennaam et al., 2014), decreased generation intervals (Schaeffer, 2006), increased rates of improvement for total merit index (TMI; e.g., García-Ruiz et al., 2016; Doublet et al., 2019), but also variable changes in genetic diversity among breeds (see Introduction section). Therefore, it is important to evaluate and monitor the possible impact of GS on genetic diversity, especially for local cattle breeds. In this study, we investigated changes in genetic diversity and population demographic structure in five local European cattle breeds from three countries. Hereafter, we first discuss the results of the observed changes in genetic diversity due to the implementation of GS in the five breeds. Then, we discuss the impact of management factors on the breeds’ genetic diversity and their population demographic structure. Finally, we discuss the implications of this study.

### Changes in genetic diversity after the implementation of GS

Overall, the observed changes in genetic diversity due to the implementation of GS for the five breeds analysed align with those reported in other studies. With the implementation of GS, we observed that the breeds’ TMI levels increased at a faster rate overall (results not shown), generation intervals for sires decreased (except for MRY), and changes in inbreeding and kinship rates varied (Table 2, Figure 4, Figure 7). For MRY, only two years of data were available after the implementation of GS; therefore, strong conclusions should be avoided. Nonetheless, for MRY, Δ*F_PED_*, Δ*f_PED_*, and Δ*F_ROH_* all increased after the introduction of GS in 2019 compared to the rate for the period 2013-2019, although both Δ*f_PED_* and Δ*F_ROH_* rates were smaller than those before 2013. For NRC, Δ*F_PED_*, Δ*f_PED_*, and Δ*F_ROH_* all decreased after the introduction of GS. Finally, after the introduction of GS, Δ*F_PED_*, Δ*f_PED_*, and Δ*F_ROH_* all decreased for ABO and VOS, while they all increased for TAR. Therefore, changes in genetic diversity after the introduction of GS were not consistent across breeds, with the timing of these changes not always corresponding to the introduction of GS. Thus, there are likely other genetic management factors influencing inbreeding and kinship rates in these breeds than just the introduction of GS *per se*.

It is not fully clear why inbreeding rates per generation (Δ*F*) increased after the introduction of GS in the major dairy cattle breeds, but not in some other breeds (Doekes, 2020). One possible reason is that, although more bulls are screened for selection with the availability of genomic information, fewer bulls are marketed, leading to a higher contribution of the selected bulls to the next generation. A lack of genomic control of inbreeding (Sonesson et al., 2012) and the strategy to update the reference population (Eynard et al., 2018a) may also play a role on the estimation of genomic-based inbreeding and on the impact on genetic diversity, respectively. Finally, intense competition between AI companies to provide bulls for the market with high breeding values for TMI may have shifted the focus to short term gains, rather than increasing genetic diversity needed for long term gain, especially in major global cattle breeds (Doekes et al., 2018; Eynard et al., 2018b; Doublet et al., 2019; Doekes, 2020). In this study, we showed that changes in Δ*F* were variable, and that increases in Δ*F* following the introduction of GS are not limited to the major global cattle breeds, as Δ*F* also increased in TAR and MRY, both of which have relatively small population sizes.

Overall, changes in *Fped* and *Froh* trends were in agreement, with increments and decrements occurring in the same period. However, differences in *ΔF* tended to be stronger when based on ROHs. For example, in ABO, *ΔFped* changed from 1.19% to 0.99% while *ΔFroh* from 2.76% to 0.68%. ROHs allow capturing inbreeding arising from ancestors in more remote generations (Keller et al., 2011; Speed and Balding, 2015). In this study, we used ROHs with a minimum length of 1,000 kb, and a rough estimate of the average number of generations captured by such ROHs length is of about 50 generations. For this rough estimate, we followed Doekes et al. (2018), assuming that the length of ROHs derived from a common ancestor *G* generations ago follows an exponential distribution with mean 1/2*G* Morgans (Browning, 2008; Speed and Balding, 2015), and a mean genetic distance of 1 Morgan per 100 Mb (Ma et al., 2015) with uniform recombination rates across the genome and across sexes. Consequently, in this study, *Froh* captured inbreeding due to ancestors long before the start of the pedigrees, potentially explaining why *Froh* levels were overall higher than *Fped* ones, especially for MRY and NRC at the start in the 1970s. Moreover, lack of deep pedigree information may lead to underestimate the level of *Fped*, while it does not affect *Froh* estimates. Overall, the average number of complete generation equivalents (Doekes et al., 2018) across years was higher for NRC and MRY compared to French breeds (Supplementary Figure S2). Nonetheless, the level of *Fped* was lower than that of *Froh* for all breeds analysed, possibly reflecting that ROHs capture more ancient inbreeding than pedigree does. Finally, another reason why Δ*Froh* and Δ*F_PED_* may differ is because genotyped animals may not be a random sample of animals in the pedigree. We checked for MRY whether the *F_PED_* of genotyped animals differed from the population average and found that *F_PED_* tended to be slightly higher for genotyped animals, especially in the last decade (results not shown). On the other hand, in VOS all animals are genotyped and estimated levels of *F_ROH_* and *F_PED_* were close for this breed.

### Impact of management factors and changes in population demographic structure

In the MRY, several changes in demographic population structure and breeding occurred besides the introduction of GS. The first change is a shift from predominantly natural matings on farm by local bulls to artificial inseminations by bulls from the national breeding program. This shift largely explains the reduction in the number of male calves born per year that later become sires, beginning as early as 1975 (Figure 2). The second change is the use in the breeding nucleus of management strategies such as ensuring that progeny-tested bulls originate from different parents and the “cold sire system” (Hiemstra and de Haas, 2004) from around 2000 onwards. Initially, the effect of such strategies is only seen in the reduced kinship of the calves that later have been selected as a breeding bull (Figure 6). Since the generation interval was above 6 years at that time and cows were generally selected on farm, it took up to 2013 before the effect of these strategies in the central nucleus was seen at the population level, after which both the average population kinship and inbreeding levels declined (Figure 6). Our results overall agree with those reported by Hiemstra and de Haas (2004), who performed a pedigree-based analysis including data up to 2004. Albeit only three years of data were available since the onset of GS in 2018, estimated kinships and inbreeding levels in MRY are on the rise again. Although the number of sires used is about the same as before GS, the skewness of sires’ contributions shows an increasing trend in the last three years (Figure 3 and Figure 5). This pattern suggests a possible increasing trend in inbreeding due to a more one-sided use of bulls available through GS. Finally, we observed higher levels of kinship for calves later becoming dams compared to calves later becoming sires (kinships levels started to diverge from the 1980-1990 period onwards), indicating that, on average, selected dams are more related than selected sires. These results highlight the possibility to reduce kinship and possible loss of genetic diversity by implementing additional management strategies on the dams’ side. In NRC, OCS has been implemented in 2010. Kinships at the population level have kept increasing with a slightly higher rate following the introduction of OCS, but declined after the introduction of GS. Moreover, the number of bulls used as sires (Figure 3) increased around 2000 and from 2010 onwards, the latter due to a more complete recording of natural matings. There was no obvious explanation for the peak in recordings of natural matings around the year 2000. Finally, it is clear that since 2017 less bulls are selected as sires per year, that these bulls are younger, and that their contribution is more even (Figure 1, Figure 2, Figure 4, and Figure 5). Thus, these results suggest that the introduction of OCS in NRC has mitigated the reduction in number of used bulls after the introduction of GS. Nonetheless, despite the increased number of calves selected as sires and dams in recent years, the kinship of selected sires also increased, especially after 2017. Such observed increases in kinship may lead to an increase in inbreeding levels in future generations.

For the French breeds, the number of bulls used as sires per year increased from around 2008 onwards, thus before the introduction of GS (Figure 3). Moreover, the reduction in the sires’ generation interval after the introduction of GS corresponded with a more equal contributions of bulls, although both of these took longer in VOS compared to other French breeds (Figure 5 and Figure 4). A likely reason for these observed trends, such as the more balanced sire’s contribution, is related to management practices as specific recommendations were provided to breeding companies prior to the implementation of GS, aiming to optimize both genetic gain and the preservation of genetic diversity (see Colleau et al., 2015). In particular, at a given constant cost, GS allows to propose more bulls to the market than progeny testing did. Moreover, all these three French breeds have an active management of their genetic diversity as they are very local, with only one or two companies in charge of the breeding scheme. Such companies aim to preserve the genetic potential of their breeds as no rescue will be possible from other sources (Doublet et al., 2019). Therefore, this management relies on the use of so-called “originality indices”, such as ORIx and ISUO, which enhance the bulls’ estimated breeding value based on its genetic distance with the current reproductive population. Additionally, the lowest level of inbreeding (pedigree and genomic) observed for VOS could possibly be due to the less intense selection pressure for production traits such as milk in this breed compared to the other French breeds analysed. Finally, for the ABO breed, lost families were reintroduced in the early 2000s in the breeding scheme by using cryopreserved semen of old bulls (born about 30 years earlier) (Jacques et al., 2023). This intervention has led to an increase in genetic diversity and variance, as well as improved performances for certain traits which had declined over time such as reproduction traits (Jacques et al., 2023). This genetic management strategy is partially reflected in the *f_PED_* trend (Figure 6), which shows a higher rate of kinship at the population level in the early 2000s, followed by a reduced rate thereafter. The effect of this intervention may appear with some delay relative to the reintroduction of lost families, likely due to the time needed for the strategy to take effect.

### Implications

High inbreeding rates pose a risk for populations. According to FAO, populations with *ΔF* above 1% per generation are threatened with extinction, and those with a *ΔF* above 0.5% are vulnerable (FAO, 2000; Meuwissen and Oldenbroek, 2017). Following from these guidelines, all breeds in this study had *ΔF* below the 1% limit, although ABO (*ΔF_PED_*) and TAR (both *ΔFped* and *ΔFroh*) were very close to such limit in the past, with ABO showing values above 1% for the period 2000-2014. Furthermore, with the exception of NRC, all breeds have, or have been in the past, above the 0.5% limit for *ΔF*. Thus, the results of this study underline how all breeds have to monitor inbreeding and kinship levels, also highlighting the importance of their genetic management. Theoretically, the most efficient genetic management is the use of optimal contributions (Meuwissen, 1997). Indeed, the implementation of population management strategies following OCS-like principles, such as the “cold sire system” and ensuring that progeny-tested bulls originate from different parents in MRY, seems to have been effective in lowering kinship levels, also accompanied by a delayed reduction in inbreeding at the population level. This delay is expected because genetic management strategies like OCS specifically targets the average kinship level of animals selected for breeding rather than the kinship of individual bulls and cows being mated. Thus, OCS ensures genetic diversity in the long term, while minimising the kinship of mating pairs works (only) in the short term.

GS has accelerated cattle breeding through, for example, the reduction of the generation interval, as also observed in this study. Misztal and Lourenco (2024) suggested that because of the acceleration brought by GS there are also increased risks for health and animal welfare when negative impacts on such traits are not noticed in time and properly addressed. Our study shows that a decreased genetic diversity is not caused by the introduction of GS *per se*. Indeed, effects of other measures, such as the introduction of OCS or OCS-like principles like the cold sire system, may have a more profound effect on genetic diversity. However, because of the acceleration in breeding due to GS, adverse effects on genetic diversity can occur more quickly than before the introduction of GS due to management practices that aim to achieve higher genetic gains, such as selecting too few and/or more related bulls. Consequently, genetic diversity management is all the more important when GS is applied. Fortunately, the availability of genomic data provides also the opportunity for a more efficient management of genetic diversity (Howard et al., 2017; Doekes et al., 2018).

## CONCLUSIONS

We investigated changes in population demographic structure and genetic diversity due to the introduction of GS in five European local cattle breeds. Overall, the implementation of GS led to a reduction in sires’ generation intervals. Moreover, we observed a reduction in the number of calves that later became sires and, for the French breeds, an increase in the number of sires used in the population, likely due to GS enabling the preselection of selection candidates in addition to screening for a higher number of young bulls. After the introduction of GS, the contribution of the top ten sires was more balanced, except for MRY, in which a balanced contribution was already present. Changes in genetic diversity did occur, but were not consistent across breeds. Rates of kinship, pedigree inbreeding, and genomic inbreeding increased in some breeds (MRY and TAR) but decreased in others (NRC, ABO, and VOS).

This study suggests that increases in inbreeding rates may occur after the introduction of GS, although they may not be directly due to the introduction of GS *per se*. The effect of population management strategies, such as the introduction of the “cold sire system” in MRY or OCS in NRC may have a much stronger effect in decreasing kinship and inbreeding rates in the longer term. Therefore, after the implementation of GS, it is essential to monitor changes in both genetic diversity and population demographic structure and adapt the breed management accordingly when necessary.

## ACKNOWLEDGEMENTS

This project has received funding from the European Union’s Horizon 2020 Programme for Research & Innovation under grant agreement n°101000226. This project is part of EuroFAANG (https://eurofaang.eu).

The authors would like to thank CRV, Mathijs Van Pelt, Gerben de Jong, and Henk Geertsema for providing the dataset and early feedback on the MRY population.

---

**Supplementary Figure S1.**
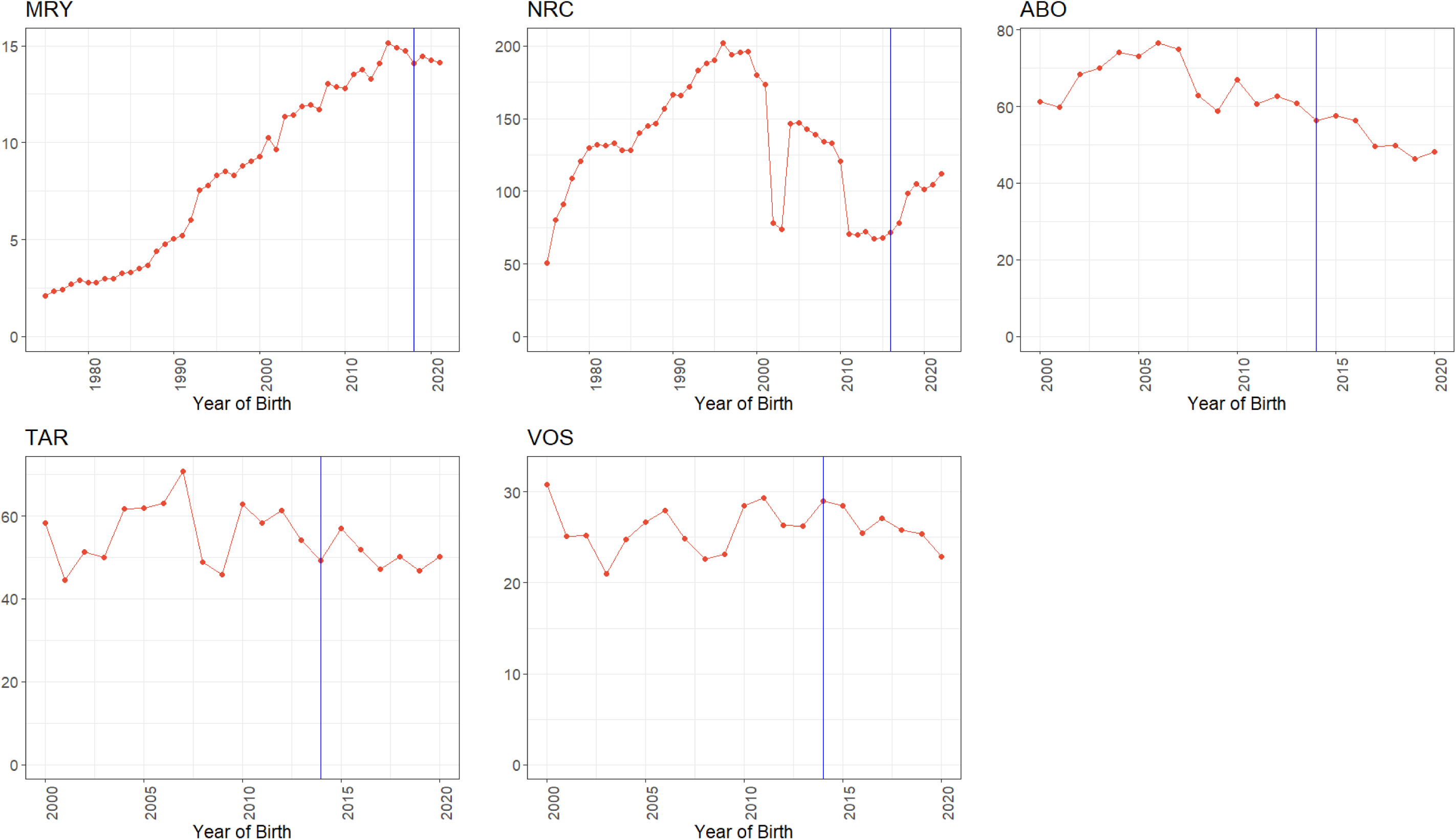
Distribution of the number of calves born per sire in each year.

**Supplementary Figure S2.**
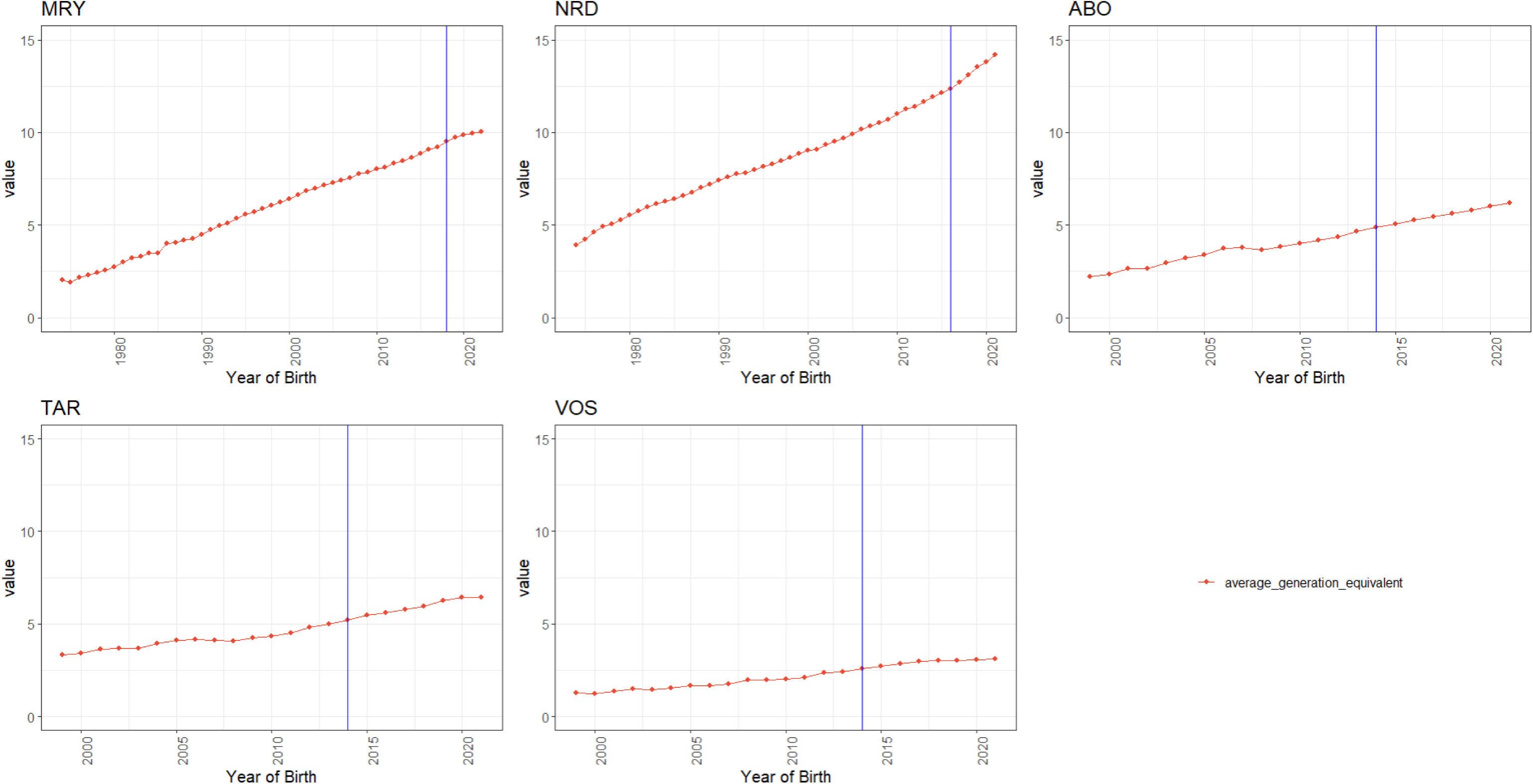
Average number of generation equivalent.

